# Single-cell proteomics maps circulating monocyte dynamics in advanced head and neck squamous cell carcinoma

**DOI:** 10.64898/2026.09.08.750082

**Authors:** Heloísa Monteiro do Amaral-Prado, Jackson Gabriel Miyamoto, Ariane Fidelis Busso-Lopes, Nilson Antônio da Rocha Coimbra, Daniella de Figueiredo, Romênia Ramos Domingues, Bianca Alves Pauletti, Ana Leticia Mores, Tiago da Silva Medina, Rodrigo Nalio Ramos, Thaís Bianca Brandão, Ana Carolina Prado-Ribeiro, Luiz Paulo Kowalski, Adriana Franco Paes Leme

## Abstract

It remains unclear whether metastatic progression in head and neck squamous cell carcinoma (HNSCC) is accompanied by functional remodeling of circulating immune cells at the proteome level. To address this question, we applied single-cell proteomics (SCP) to cryopreserved peripheral blood mononuclear cells (PBMCs) from three patients with HNSCC representing distinct stages of metastatic progression and two healthy donors. Proteomic analysis of 619 individual PBMCs resolved major immune cell populations, with up to 1,638 proteins quantified per cell. Trajectory inference, clustering, and differential abundance analyses revealed proteomic remodeling associated with disease progression, with the most pronounced changes occurring within a monocyte cluster composed exclusively of cells from patients with nodal metastasis. This population showed increased abundance of interferon-related proteins, HLA molecules, and myeloid immunoregulatory signatures, consistent with an activated, interferon-associated monocyte state. In parallel, lymphocytes showed reduced coordinated abundance of proteins associated with activation, cytotoxicity, and degranulation, consistent with altered cytotoxic effector programs across disease stages. Overall, these findings reveal distinct proteomic states in circulating monocytes and lymphocytes associated with HNSCC progression and identify selective remodeling of the monocyte compartment in nodal metastatic disease.

## Introduction

Head and neck squamous cell carcinoma (HNSCC), the predominant subtype of head and neck cancers (HNC), is characterized by a high global incidence and poor prognosis in advanced disease, particularly in the presence of lymph node metastasis, which reduces the 5-year survival rate to below 50% ^1–3^. Although significant efforts have focused on understanding immune-cell interactions within the tumor microenvironment, metastatic progression is increasingly recognized as a systemic process involving dynamic remodeling of the peripheral immune system ^4,5^. Because circulating immune cells continuously integrate tumor-derived and host immune signals, peripheral blood is an accessible, minimally invasive liquid biopsy that provides a real-time snapshot of the systemic immune landscape, enabling monitoring of immune remodeling throughout disease progression ^5,6^.

Previous bulk proteomic studies have identified circulating protein signatures associated with lymph node metastasis in HNSCC, suggesting altered immune responses primarily involving myeloid cells and lymphocytes ^7,8^. However, because bulk proteomics averages signals across heterogeneous immune-cell populations, the cellular origin and functional organization of these proteomic alterations remain understudied. Therefore, profiling these cells by mass spectrometry-based single-cell proteomics (SCP) enables the characterization of cellular functional heterogeneity associated with disease progression ^9,10^.

Recent single-cell transcriptomic sequencing (scRNA-seq) studies have comprehensively characterized immune-cell populations within the HNSCC tumor microenvironment, revealing dynamic remodeling of both myeloid and lymphoid compartments during disease progression ^11,12^. Nevertheless, whether these transcriptional programs are accompanied by coordinated remodeling of the circulating proteome remains largely unknown. As the primary executors of cellular function, proteins provide direct information on signaling pathways, immune activation, metabolic adaptation, and effector programs that cannot be fully inferred from transcript abundance alone. Despite significant technological advances that have remarkedly increased proteome coverage, the translation of SCP to clinically relevant patient cohorts remains challenging because of limited access to well-characterized clinical samples, low analytical throughput, and the technical complexity of processing primary human samples ^13–18^. Consequently, most published SCP studies with patient-derived samples have analyzed relatively small cohorts, highlighting the emerging nature of clinical SCP applications ^13,14^.

Here, we applied a label-free SCP workflow to 619 cryopreserved peripheral blood mononuclear cells (PBMCs), without previous cell sorting, from patients with HNSCC representing distinct stages of nodal metastatic progression and healthy donors, with the hypothesis that circulating immune cells undergo proteomic remodeling during disease progression. Up to 1,638 proteins were quantified per cell, enabling the identification of major immune cell populations and revealing proteomic remodeling associated with disease progression predominantly within a monocyte cluster exclusively in patients with nodal metastasis. By comparing our SCP data with an independent scRNA-seq dataset from primary HNSCC tumors ^11^, we identified convergent interferon-associated and immunoregulatory programs in the myeloid compartment across protein and transcriptomic measurements. Collectively, these orthogonal observations highlight complementary information provided by SCP and scRNA-seq, with SCP resolving protein-level immune states in circulating cells that cannot be directly inferred from transcript abundance alone.

Collectively, beyond validating transcriptomic observations, SCP resolved dynamic monocyte and lymphocyte states linked to nodal metastasis that cannot be inferred from transcript abundance alone, establishing single-cell proteome-level immune phenotyping as an important strategy for investigating systemic cancer immunity.

## Results

### From the bedside to the bench: Quality controls and metrics of SCP data generated

Blood samples from patients representing progressive stages of HNSCC and healthy donors (denominated as control_1 and control_2) were collected at the hospital for SCP analysis (**Figure 1A**). The HNSCC cases selected for this analysis covered the progression of disease severity as defined by the tumor size (T), lymph node involvement (N), and metastasis (M) (TNM) staging system ^19^. However, this system has several limitations, as patients with the same TNM stage often exhibit distinct clinical behaviors, differential treatment responses, and substantial variability in clinical outcomes ^20^. The patients analyzed in this study are classified as pT4aN0 (pN0), which represented a large, locally invasive tumor that had extended into deep anatomical structures but showed no pathological evidence of regional lymph node metastasis, and pT2N2b (pN+), which represented a medium-sized primary tumor accompanied by multiple ipsilateral lymph node metastases, reflecting more advanced regional dissemination. The most severe case, pT4aN3b (pN+), had extensive local invasion combined with lymph node metastasis with extranodal extension, a high-risk feature associated with aggressive biological behavior and poor clinical outcomes. None of the patients had distant metastasis (M0).

**Figure 1.**
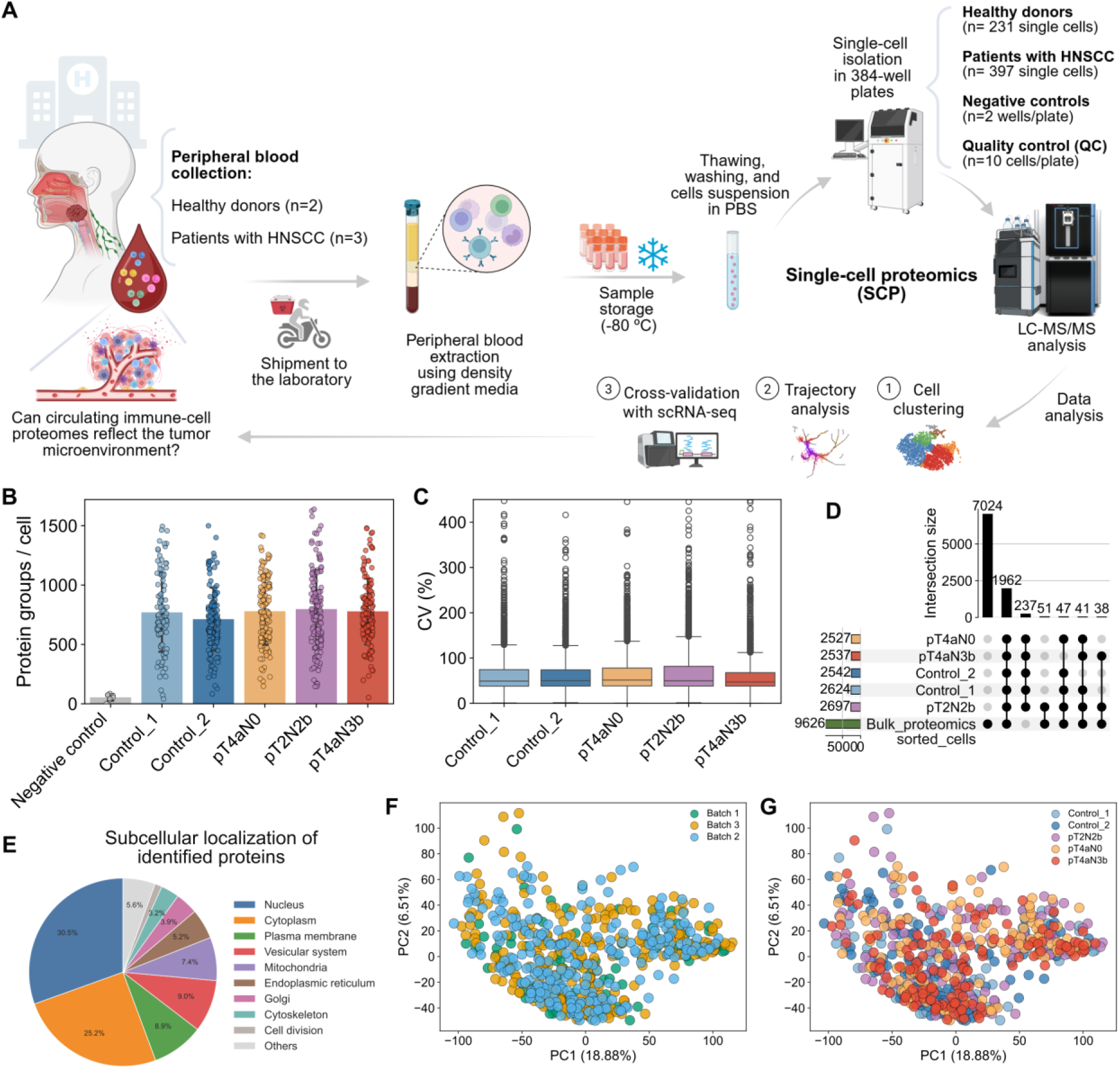
Experimental design and quality assessment of SCP of PBMCs from controls and patients with HNSCC. **(A)** Schematic overview of the experimental workflow. **(B)** Average ± standard deviation (SD) of PGs (Negative control – n = 8 wells; Control 1, n = 98 cells; Control 2, n = 133 cells; pT4aN0, n = 129 cells; pT2N2b, n = 134 cells; pT4aN3b, n = 134 cells). **(C)** Distribution of the coefficient of variation (CV) of quantified protein abundances for each experimental group, demonstrating comparable quantitative reproducibility across donors and patients. **(D)** UpSet plot showing the overlap of identified protein groups among the five biological samples, and with bulk PBMC proteomics from Rieckmann *et al.* (2017). Most of the proteins identified by SCP overlapped with previously reported PBMC bulk proteomes. **(E)** Subcellular localization of all identified proteins in sPBMCs (n=3,118 protein groups). Human Protein Atlas database was used as a reference for subcellular protein localization. A total of 2,557 proteins identified in our SCP dataset were shared with the database. Principal component analysis (PCA) colored according to LC-MS/MS acquisition batch **(F)** and clinical groups **(G)**, showing no evident batch-dependent segregation.

Together, these stages captured a gradient of tumor progression, from locally advanced disease to extensive regional spread. Cryopreserved single PBMCs (sPBMCs) were isolated in 384-well plates as single cells using the cellenONE X1 platform and analyzed by data-independent acquisition (DIA) liquid chromatography–tandem mass spectrometry (LC–MS/MS) on an Orbitrap Astral mass spectrometer (**Figure 1A**).

To ensure analytical robustness, all acquisition plates included cryopreserved single HeLa cells (n=10) as internal quality controls (QC) (**Figure S1A**). Peptides and protein groups (PGs) identifications remained highly consistent across plates, with approximately 5% variation throughout the multi-day experiment (**Figures S1B, C**; **Table S1**). In parallel, Principal Component Analysis (PCA) and instrument QC metrics further confirmed the absence of relevant plate-to-plate variability and the overall stability of the LC–MS/MS platform throughout data acquisition (**Figures S1D, E**). Blank wells (denominated as negative controls; n=8) were also included to identify background proteins, which were subsequently excluded from downstream analyses (**Figure 1B**, **Table S2**). sPBMCs yielded approximately 800 quantified PGs per cell on average, with up to 1,638 PGs identified in individual cells (**Figure 1B**, **Table S3**). Notably, our workflow achieved this depth directly from heterogeneous PBMC samples, without prior isolation or enrichment of specific immune cell populations ^13,18,21,22^, highlighting its suitability for unbiased profiling of clinically relevant samples.

Moreover, quantitative reproducibility was comparable across all conditions, with no significant differences in coefficients of variation (CVs) between clinical groups (**Figure 1C**). These quality-control analyses demonstrate that the SCP workflow generated reproducible and technically robust single-cell proteomes, providing a reliable foundation for investigating whether metastatic progression is accompanied by remodeling of circulating immune-cell states. Also, more than 3,100 PGs were identified across the entire cohort, demonstrating broad proteome coverage despite the intrinsic heterogeneity of circulating immune cells. Across all patient groups, 2,199 PGs were consistently detected in every experimental group, representing a robust core proteome (**Figure 1D**). Comparison with a previously published bulk-sorting proteomics dataset based on PBMC subpopulations ^23^ revealed that 2,683 PGs overlapped between the datasets, with 1,962 PGs shared in all patient groups (**Figure 1D**). Subcellular localization analysis of the identified proteins using the Human Protein Atlas showed that this shared proteome was broadly distributed across major cellular compartments, with proteins predominantly localized to the nucleus (30.5%) and cytoplasm (25.2%), followed by the plasma membrane (8.9%), vesicular system (9.0%), mitochondria (7.4%), endoplasmic reticulum (5.2%), and Golgi apparatus (3.9%), indicating comprehensive coverage of proteins involved in diverse cellular structures and biological functions (**Figure 1E**).

Next, because samples were acquired across multiple analytical plates (**Figure S1A**) and, consequently, on different experimental days, we evaluated whether batch effects contributed to the observed variability. PCA showed no segregation according to the acquisition plates, with the first two principal components (PC1, 18.88% and PC2, 6.51%) explaining 25.39% of the total variance (**Figures 1F, G**). In addition, a significant positive correlation between cell diameter (µm) and the number of identified PGs was consistently observed across all groups (Spearman’s ρ = 0.5–0.7, p-value < 0.05), supporting cell size as an important determinant of proteome depth ^24,25^ (**Figure S2**, **Table S4**).

Together, these analyses demonstrate that the SCP workflow generated technically robust and comprehensive proteome profiles of circulating immune cells, with a proteome depth comparable to that achieved in recent state-of-the-art single-cell proteomics studies of primary immune cells ^13,18,21,22^, while avoiding prior cell sorting or lineage enrichment, making it well suited for downstream characterization of disease-associated immune remodeling.

### Single-cell proteomics resolves the major circulating immune populations in HNSCC

To characterize the cellular composition of the SCP dataset, we performed unsupervised clustering of circulating immune-cell proteomes. Uniform Manifold Approximation and Projection (UMAP) identified multiple well-resolved Leiden clusters, indicating proteomic heterogeneity among circulating immune cells (**Figure 2A**; **Table S5**). All clinical groups contributed to the major cellular clusters (**Figure 2B**), suggesting that the observed structure primarily reflected biological heterogeneity rather than technical segregation. Nevertheless, differences in cluster occupancy across samples suggested progressive disease-associated remodeling of specific immune populations (**Figure 2C**).

**Figure 2.**
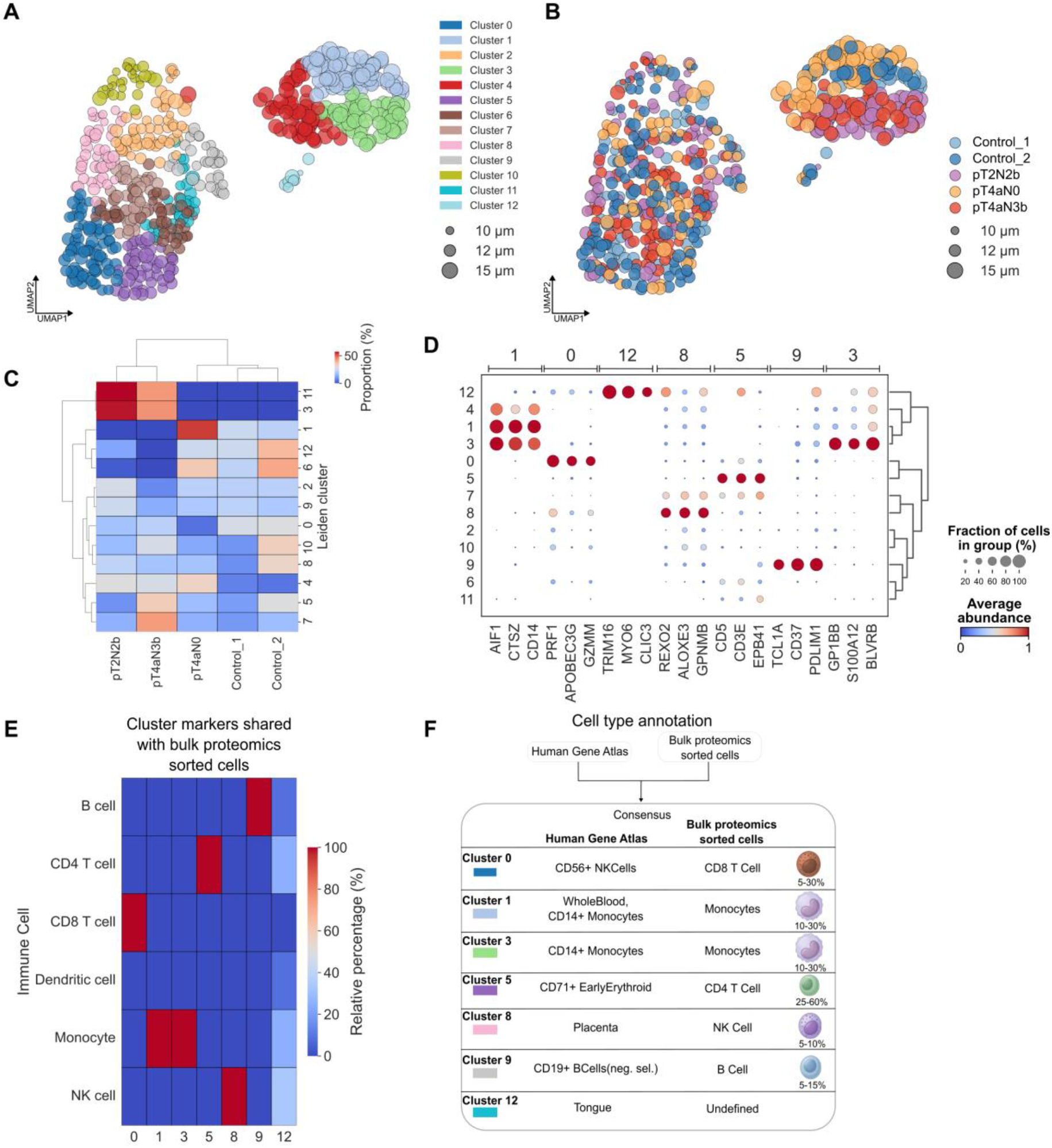
Single-cell proteomics reveals immune cell heterogeneity and patient-specific remodeling in circulating PBMCs from HNSCC. **(A)** UMAP representation of 619 sPBMCs colored by Leiden clusters (resolution of 1.5). Dot size reflects the estimated cell diameter (10–15 µm). **(B)** UMAP colored by sample origin, including healthy controls and HNSCC patients at different clinical stages (Control_1, Control_2, pT4aN0, pT2N2b, and pT4aN3b). **(C)** Heatmap showing the relative abundance of cells from each condition across Leiden clusters. Colors represent the percentage of cells contributed by each condition within each cluster. Hierarchical clustering (method=“ward”, metric=“euclidean”) was applied to both rows and columns to identify similarities in cluster composition. **(D)** Dot plot of representative cluster-specific protein markers identified using the Wilcoxon rank-sum test. Dot size indicates the fraction of cells expressing each protein within a cluster, whereas color intensity represents the average normalized protein abundance. **(E)** Heatmap showing the overlap between cluster-specific protein markers identified by SCP and immune cell markers derived from a reference bulk proteomics dataset generated from sorted human immune cell populations. Values represent the relative percentage of shared markers assigned to each immune cell type within each Leiden cluster. **(F)** Consensus annotation of Leiden clusters obtained by integrating protein signatures with the Human Gene Atlas and canonical bulk proteomics-sorted immune cell markers. The table summarizes the inferred immune cell identity for each cluster, together with the estimated cell proportion expected of total isolated PBMCs.

Since cell size is an important morphological feature that differs across PBMC subsets and influences proteome depth ^24,25^, we next examined whether cellular morphology contributed to the observed clustering patterns. Overall, PBMC populations differ markedly in cell size, including small naïve lymphocytes (∼6–8 µm in diameter), activated lymphocytes (∼10–15 µm), and larger cell types such as monocytes and dendritic cells (∼15–20 µm) ^15,26,27^. As expected, larger cells (∼14 μm), corresponding predominantly to monocyte-associated clusters, segregated from smaller lymphocyte-enriched populations (10–13 μm), consistent with the known morphological diversity of PBMCs (**Figures 2A, B**). Although all clinical groups contributed to the major cellular clusters, their relative representation differed markedly across conditions (**Figure 2C**). Notably, the two controls exhibited highly similar cluster occupancy patterns, supporting the reproducibility of the SCP workflow. In contrast, the pN0 group showed preferential enrichment in cluster 4, whereas the metastatic patients (pN+) displayed marked enrichment in clusters 3 and 11, together with increased representation in cluster 7 for the most advanced case (pT4aN3b). These progressive shifts in cluster occupancy suggest that HNSCC progression is accompanied by extensive remodeling of the circulating immune compartment, reflecting transitions between proteomic cell states within established immune populations rather than the emergence of new immune-cell populations.

To further characterize the immune cell populations derived from this SCP dataset, Leiden cluster markers (**Figure 2D**, **Table S6**) were then intersected against a reference bulk proteomics dataset generated from sorted human immune cell populations ^23^ **(Figure 2E**), as well as with the Human Gene Atlas (HGA) database, to reach a consensus on the probable cell type of each cluster (**Figure 2F**). This strategy consistently identified the major PBMC populations, including monocytes, CD8^+^ T cells, CD4^+^ T cells, NK cells and B cells. Monocyte identity was supported by canonical proteins including CD14, CYBA and CYBB (Clusters 1 and 3), whereas CD8^+^ T cells were characterized by PRF1 and GZMM (Cluster 0), CD4^+^ T cells by CD3E and CD5 (Cluster 5), NK cells by REXO2 and GPNMB (Cluster 8), and B cells by TCL1A and CD37 (Cluster 9). Although cluster 4 clearly displayed a monocyte proteomic signature, no uniquely enriched protein markers were identified to distinguish it from the other monocyte clusters, precluding its annotation as a separate monocyte subpopulation. In contrast, cluster 12 remained undetermined because its protein signature showed limited similarity to established immune cell reference profiles. Also, the small number of cells in this cluster further limited confidence in its annotation. Overall, the concordance between cluster-specific protein markers and bulk proteomics-derived immune signatures supported the robustness of the cell-type annotations obtained by SCP. In comparison, HGA annotations were less consistent for some clusters (8 and 12), suggesting that the bulk proteomics reference better captured the immune cell identities represented in the SCP dataset.

Finally, these analyses demonstrate that the untargeted SCP workflow resolved the major circulating immune populations present in PBMCs. More importantly, differences in cluster occupancy across clinical groups suggested progressive remodeling of specific immune populationsImmune composition is remodeled during metastatic progression

We next investigated how immune cell composition changed throughout HNSCC progression (**Figures 3, 4**). Monocyte-associated clusters were identified by CD14 protein abundance enrichment and annotated according to cell population definitions established by reference bulk-sorting proteomics, whereas lymphocyte-associated clusters were identified by the enrichment abundance of PTPRCAP protein (also known as CD45-associated protein (CD45-AP)) and the corresponding reference proteomic signatures (**Figure 3A**). This classification revealed a progressive shift in immune composition. Controls were predominantly enriched in lymphocyte-associated populations, while patients with HNSCC exhibited expansion of monocyte-associated cells accompanied by a relative reduction in lymphocytes (**Figure 3B**). These findings are consistent with previous clinical studies demonstrating that a reduced lymphocyte-to-monocyte ratio is associated with poor prognosis and decreased overall survival in patients with HNSCC ^28,29^.

**Figure 3.**
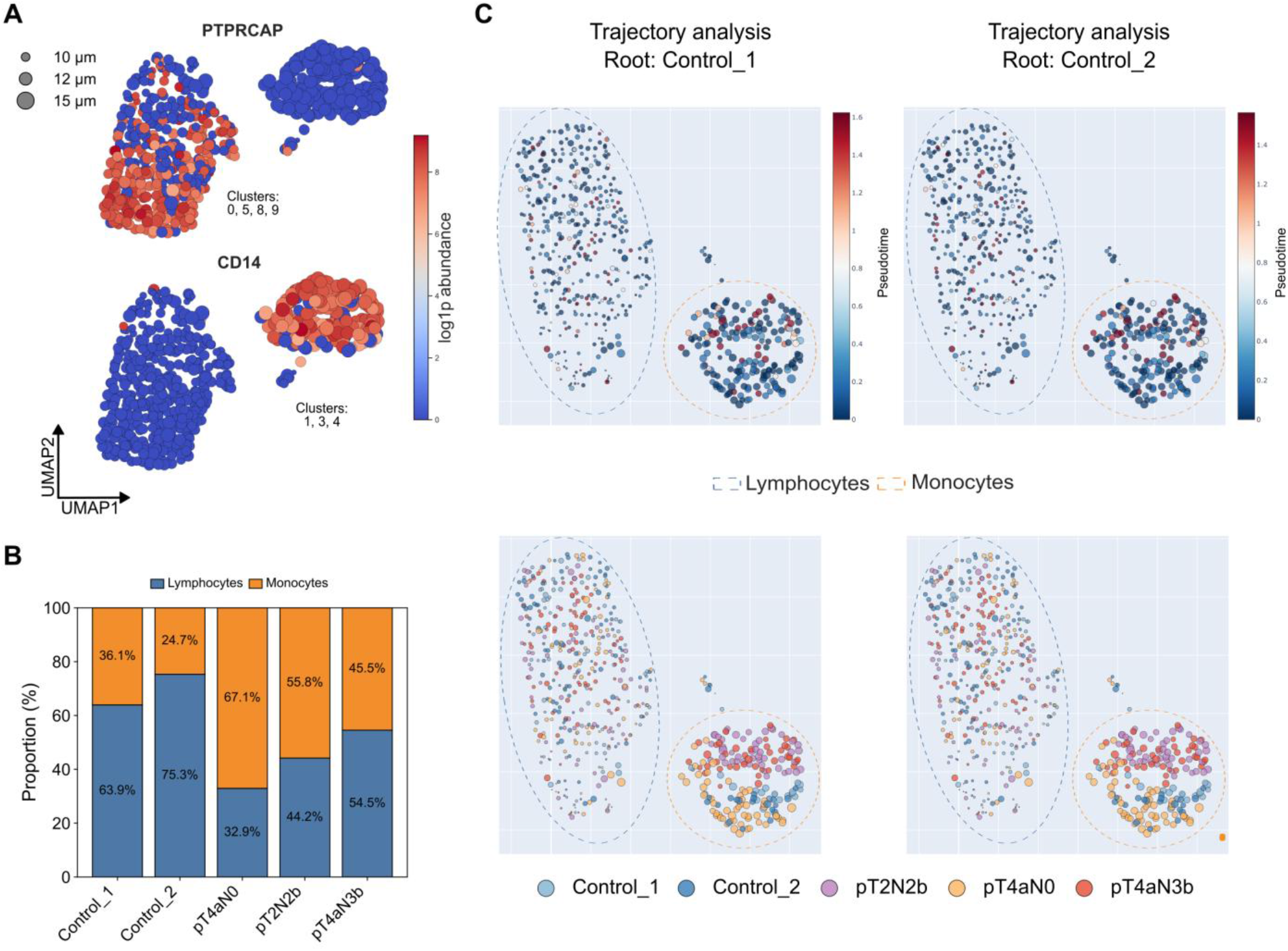
Cell-type composition and pseudotime trajectory analysis of sPBMCs from HNSCC patients. **(A)** UMAP visualization of representative marker proteins highlighting the distribution of lymphocyte (PTPRCAP) and monocyte (CD14) populations. Dot size represents cell diameter, and color intensity indicates log2-transformed protein abundance. **(B)** Relative proportions of lymphocytes and monocytes across controls and HNSCC patients. Cell identities were assigned based on the Leiden clusters highlighted in **(A)**. A total of 619 cells were analyzed, including 211 lymphocytes, 186 monocytes, and 222 other cells. **(C)** Pseudotime trajectory analysis inferred using Monocle3 with the Scanpy-derived UMAP embedding. To evaluate the robustness of trajectory inference, pseudotime was independently reconstructed using Control_1 or Control_2 as the root state. UMAPs display the inferred pseudotime and clinical groups overlaid on the cellular embedding. Dot sizes are in accordance with the cell diameter.

Changes in immune-cell abundance alone cannot distinguish whether disease progression results from the expansion of discrete cell populations or from gradual transitions between cellular states ^30–32^. We therefore additionally performed an exploratory trajectory analysis to reconstruct the continuous proteomic remodeling associated with HNSCC progression. To evaluate the robustness of trajectory inference, pseudotime was reconstructed twice, independently using each control as the root state (pseudotime = 0), representing the earliest immune proteomic state from which disease-associated transitions were inferred (**Figure 3C**; **Figure S3**). Trajectory inference reconstructed highly similar topologies regardless of which control was used as the root state, demonstrating that the inferred immune-state transitions were robust to root selection (**Figure S3A**). In both reconstructions, cells from patients with HNSCC extended toward progressively later pseudotime states relative to controls, indicating continuous disease-associated immune-state remodeling (**Figure S3B**). Importantly, a highly similar trajectory was reconstructed when the pN0 patient was used as the root, indicating that the overall trajectory topology is robust to the choice of the starting population and is primarily determined by the underlying single-cell proteomic information (**Figure S3B**).

### Trajectory analysis reveals a conserved proteomic signature of metastatic progression

A total of 422 proteins were consistently associated with the inferred trajectory regardless of the selected control, demonstrating the robustness of the reconstructed pseudotime. Functional enrichment analysis revealed that these proteins were predominantly involved in vesicle-mediated transport, intracellular trafficking, and related membrane dynamics, suggesting that progressive immune remodeling is accompanied by coordinated changes in protein transport and vesicular processes (**Figure 4A**).

**Figure 4.**
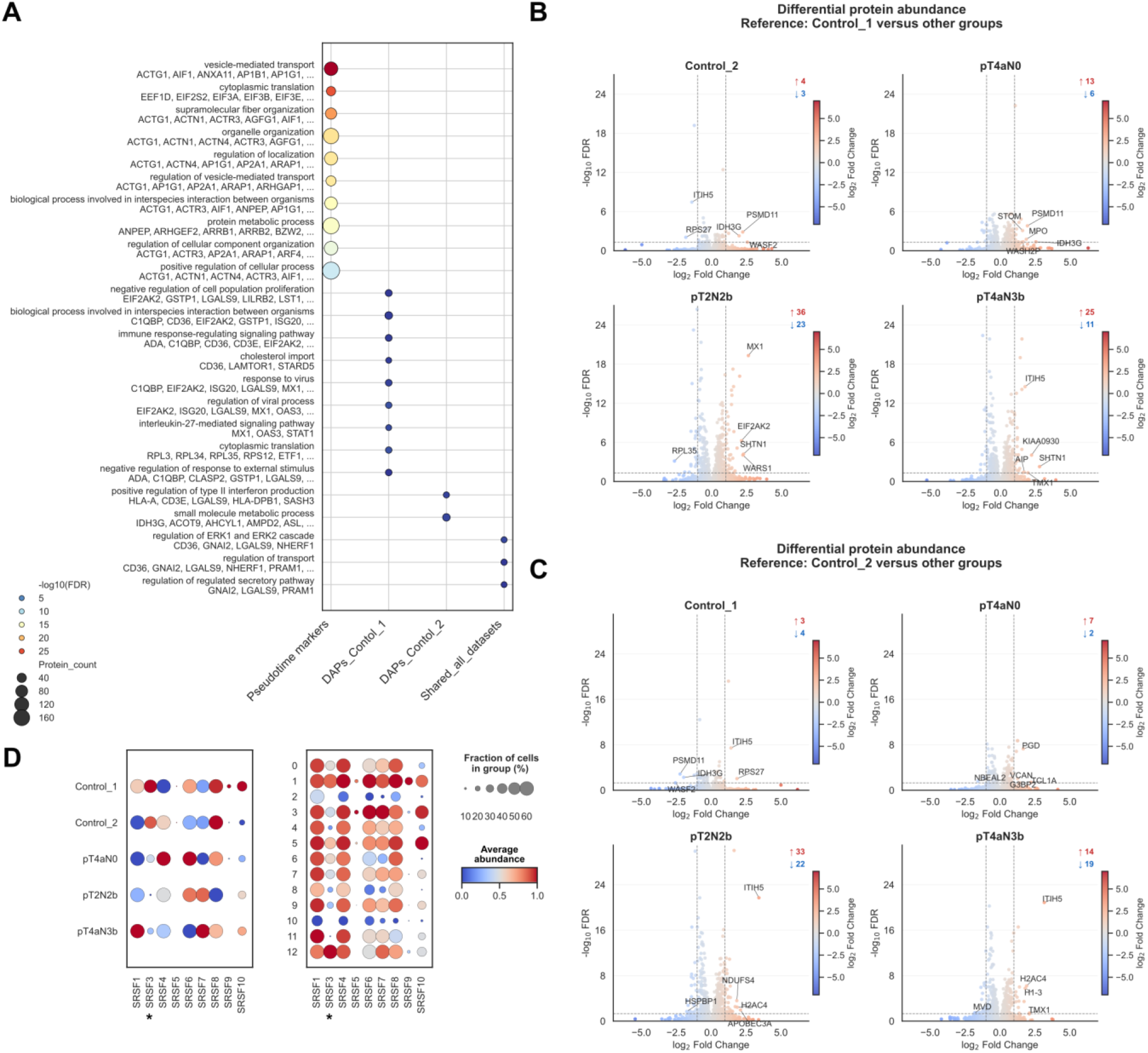
Trajectory-associated proteins and differential protein abundance during HNSCC progression. **(A)** Gene Ontology (GO) biological process enrichment analysis of proteins associated with pseudotime progression and differentially abundant proteins (DAPs). The left panel shows the biological processes significantly enriched among pseudotime-associated proteins identified by trajectory inference. The middle and right panels summarize the enriched biological processes among DAPs identified using Control_1 or Control_2 as the reference, respectively. Dot size represents the number of proteins associated with each GO term, and dot color indicates enrichment significance (−log10 adjusted P value). **(B–C)** Volcano plots showing differential protein abundance across experimental groups using Control_1 **(B)** or Control_2 **(C)** as the reference. Comparisons were performed between the two controls, as well as the controls and HNSCC patients (pT4aN0, pT2N2b, and pT4aN3b). The top five proteins satisfying the significance thresholds (adjusted p < 0.05 and |log2 fold change| ≥ 1) are highlighted. **(D)** Dot plots of the average abundance of splicing-related proteins that were differentially abundant in OSCC patients when compared with Control_1 or Control_2. Dot size indicates the fraction of cells expressing each protein within a patient group or Leiden cluster, whereas color intensity represents the average normalized protein abundance. * Splicing-related protein with abundance in accordance with bulk proteomics from Busso-Lopes et al. ^8^

To identify disease-associated proteomic alterations independently of donor-specific effects, differential abundance analyses were performed using each control as an independent reference (**Figures 4B, C**). In both analyses, the number of differentially abundant proteins (DAPs) increased progressively with disease severity, with the largest differences observed between controls and metastatic patients (**Figures 4B, C**; **Table S8**). In contrast, relatively few DAPs distinguished the non-metastatic patient from controls, indicating that the most extensive proteomic remodeling accompanies metastatic progression rather than primary tumor burden.

Intersecting trajectory-associated proteins with DAPs identified in both control-based analyses revealed a set of ten proteins (BLVRB, CD36, GNAI2, LAP3, LGALS9, NHERF1, PRAM1, TMX1, TYMP, and VASP). These proteins are described as participating in vesicle-mediated trafficking, cytoskeletal organization, myeloid-cell activation, and immune regulation, reinforcing the biological processes highlighted by the trajectory analysis (**Figure 4A**). Notably, LGALS9 has established roles in monocyte activation and inflammatory signaling, further supporting monocyte remodeling as a central feature of HNSCC progression ^33,34^.

Remarkably, although splicing-related proteins were not among the trajectory-associated markers, several members of the Serine/Arginine-rich Splicing Factor family (SRSF) exhibited consistent abundance changes across HNSCC patients (**Figure 4D**). SRSF3, SRSF4, and SRSF8 were decreased in pN+ patients, whereas SRSF7 was increased. These alterations were broadly distributed across immune populations and were consistent with our previous bulk proteomics study, which also identified reduced SRSF3 abundance in HNSCC-associated immune compartments ^8^. Together, these findings suggest that dysregulation of RNA-splicing machinery accompanies systemic immune remodeling during disease progression.

### Monocyte subset undergoes functional reprogramming during metastatic progression

Although both lymphoid and monocyte populations extended toward later pseudotime states during disease progression, several observations indicated more pronounced remodeling in the monocyte compartment. Monocyte-associated populations expanded in patients with HNSCC, and cluster 3 was composed exclusively of cells from pN+ patients (**Figure 2C**), suggesting the emergence of a disease-associated monocyte state. This pattern was further supported by unsupervised hierarchical clustering of the SCP data, in which cluster 3 formed a distinct pattern characterized by coordinated changes in protein abundance and cells occupying late pseudotime (**Figure S4**). Therefore, because it well known that activated monocytes undergo coordinated changes in inflammatory and interferon signaling, immune regulation, and oxidative stress ^35^, we next investigated the abundance of representative proteins associated with these processes across the Leiden clusters (0–12) and clinical groups (**Figure S5**) to determine whether the metastatic-associated monocyte population exhibited an activated phenotype.

Overall, the pseudotime reconstruction, unsupervised proteomic clustering, and protein signature abundance patterns by clinical groups and clusters consistently identify the emergence of a distinct monocyte state in the pN+ patients. This population was characterized by coordinated activation of interferon signaling, immune regulatory pathways, and cellular stress-response programs, indicating that disease progression is accompanied by selective remodeling of the monocyte compartment rather than uniform proteomic changes across all immune cell populations (**Figure S5**).

### Monocyte subclustering reveals heterogeneous subset-associated proteomic states

To further investigate heterogeneity within the monocyte compartment, we performed a subclustering analysis restricted to monocyte-associated cells (clusters 1, 3, and 4 from **Figure 2**) (**Figure 5**). We identified six monocyte subclusters with distinct distributions across clinical groups (**Figures 5A, B, H**), as well as distinct patterns of HLA-associated, myeloid suppressive, myeloid-associated, and interferon-stimulated proteins (**Figure 5D–G**), indicating that the monocyte compartment comprises multiple distinct states at the protein level.

**Figure 5.**
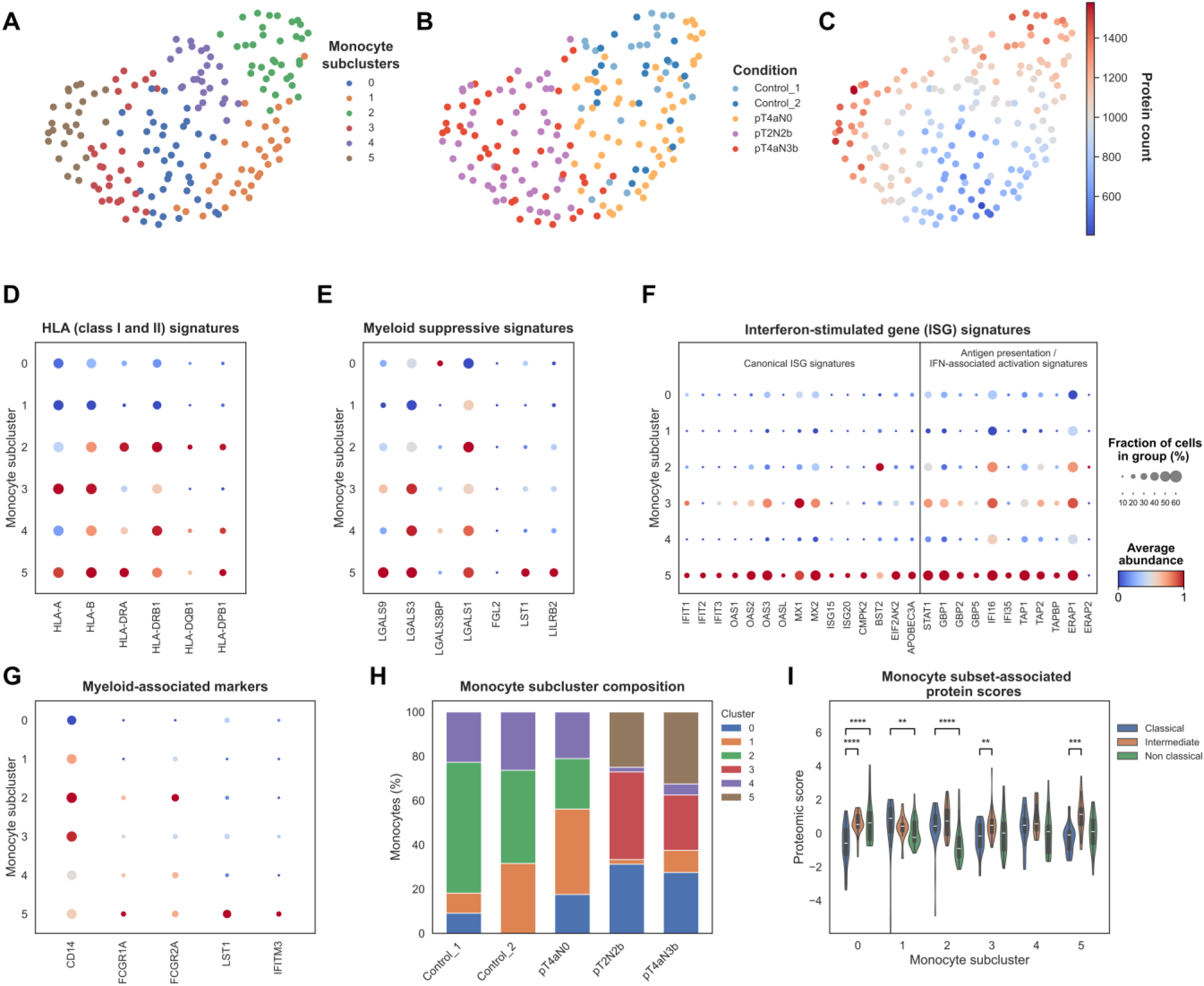
Monocyte subclustering reveals heterogeneous proteomic states associated with classical, intermediate, and nonclassical monocyte programs. **(A)** UMAP representation of monocyte subclusters identified by Leiden clustering (resolution of 1). **(B)** UMAP representation colored according to experimental condition. **(C)** Distribution of the number of quantified proteins across individual monocytes. **(D–G)** Dot plots showing the abundance patterns of selected protein signatures across monocyte subclusters, including HLA class I and II signatures **(D)**, myeloid suppressive signatures **(E)**, interferon-stimulated gene (ISG) signatures, comprising canonical ISGs and antigen presentation/IFN-associated activation signatures **(F)**, and myeloid-associated markers **(G).** Dot size represents the percentage of cells in each subcluster in which the protein was detected, while dot color represents the relative protein abundance pattern. **(H)** Relative composition of monocyte subclusters across the indicated conditions. **(I)** Violin plots showing protein scores associated with classical (CD14, S100A8, S100A9, FCN1, LYZ, VCAN, CTSZ, MNDA, S100A10), intermediate (HLA-DRA, HLA-DPA1, HLA-DPB1, CD74, CTSS, FCGR1A, LILRB1, LGALS3, IFITM3), and nonclassical (LRP1, LST1, LILRB1, SERPINA1, CEACAM8, MMP8) monocyte subsets across monocyte subclusters. Paired Wilcoxon signed-rank tests were performed within each monocyte subcluster, using the classical monocyte-associated protein score as the reference and comparing it with the intermediate- and nonclassical-associated scores. Brackets indicate significant comparisons, with \**p* < 0.01, \*\**p* < 0.001, \*\*\**p* < 0.0001, and \*\*\*\**p*<0.00001. Comparisons not reaching statistical significance are not shown.

Notably, the distribution of monocyte subclusters differed across clinical groups (**Figure 5H**). While subcluster 2 was predominant in controls and pT4aN0, subclusters 3 and 5 presented a larger proportion of monocytes in the pN+ groups. Also, these subclusters displayed different molecular features, with subcluster 3 showing increased HLA- and ISG-associated proteins, whereas subcluster 5 exhibited a prominent ISG-associated profile (**Figure 5D–G**). These observations suggest that disease progression may be accompanied not only by expansion of the monocyte compartment but also by changes in the relative abundance and proteomic state of specific monocyte subpopulations

Next, we performed an exploratory analysis to check whether these proteomic states reflected features associated with canonical monocyte subsets. We calculated protein scores representing classical, intermediate, and nonclassical monocyte-associated programs (**Figure 5I**). Importantly, because SCP detects a limited number of proteins per cell and canonical monocyte subset markers were not consistently detected across cells, individual cells were not assigned to discrete monocyte subsets based on single-marker abundance. Instead, scores were calculated based on the relative abundance of proteins previously associated in the literature and the Human Protein Atlas database with classical (CD14, S100A8, S100A9, FCN1, LYZ, VCAN, CTSZ, MNDA, S100A10), intermediate (HLA-DRA, HLA-DPA1, HLA-DPB1, CD74, CTSS, FCGR1A, LILRB1, LGALS3, IFITM3), and nonclassical (LRP1, LST1, LILRB1, SERPINA1, MMP8, CEACAM8) monocytes ^36–39^.

Interestingly, the scores revealed distinct patterns of subset-associated protein programs across monocyte subclusters (**Figure 5I**; **Figure S6**). Subcluster 0 (predominant in pN+) showed higher intermediate- and nonclassical-associated scores than the classical score, whereas subclusters 3 and 5 (also predominant in pN+) showed distinct subset-associated proteomic patterns, with both exhibiting significantly higher intermediate monocyte-associated scores relative to the classical score. Subcluster 2 (most predominant in controls followed by pN0) also showed a higher intermediate-associated score relative to the classical score. In contrast, subcluster 4 (predominant in controls and pN0) did not show significant differences among the evaluated subset-associated scores. Finally, cluster 1 (predominant in Control_1 and pN0), showed a higher classical-associated score than the nonclassical-associated score, with no significant difference between classical and intermediate-associated scores.

Overall, these patterns observed suggest a tendency toward intermediate- and nonclassical-associated monocyte programs in nodal metastatic patients, whereas controls and pN0 displayed a more homogeneous distribution across the three subset-associated programs. Nevertheless, these results should be interpreted as exploratory, given the limited proteomic depth and incomplete detection of canonical markers in SCP, such as CD16 protein for nonclassical monocytes.

### Independent scRNA-seq analysis reveals a convergent interferon-associated myeloid program

To assess whether a convergent myeloid program could be identified in an independent molecular layer, we analyzed a publicly available scRNA-seq dataset from a large cohort of primary tumors from patients with HNSCC (6 pN+ and 7 pN0) and controls (n=6) generated by Choi et al. (2023) ^11^ (**Figure 6**). Tumor-associated monocytes and macrophages exhibited marked enrichment of ISGs, antigen-presentation pathways, and myeloid suppressive signatures (**Figure 6**, **Table S9**). These transcriptional programs followed a progressive pattern, increasing from controls to pN0 tumors and reaching their highest levels in pN+ cases. Although the transcriptomic dataset and our SCP cohort represent independent patient populations and distinct biological compartments (immune cells from primary tumor tissues versus circulating immune cells from peripheral blood), the observation of convergent interferon-associated myeloid programs across these independent cohorts supports the biological relevance of this immune-state signature. However, because these comparisons were performed across independent cohorts and analytical platforms, they should be interpreted as complementary and exploratory rather than direct quantitative validations.

**Figure 6.**
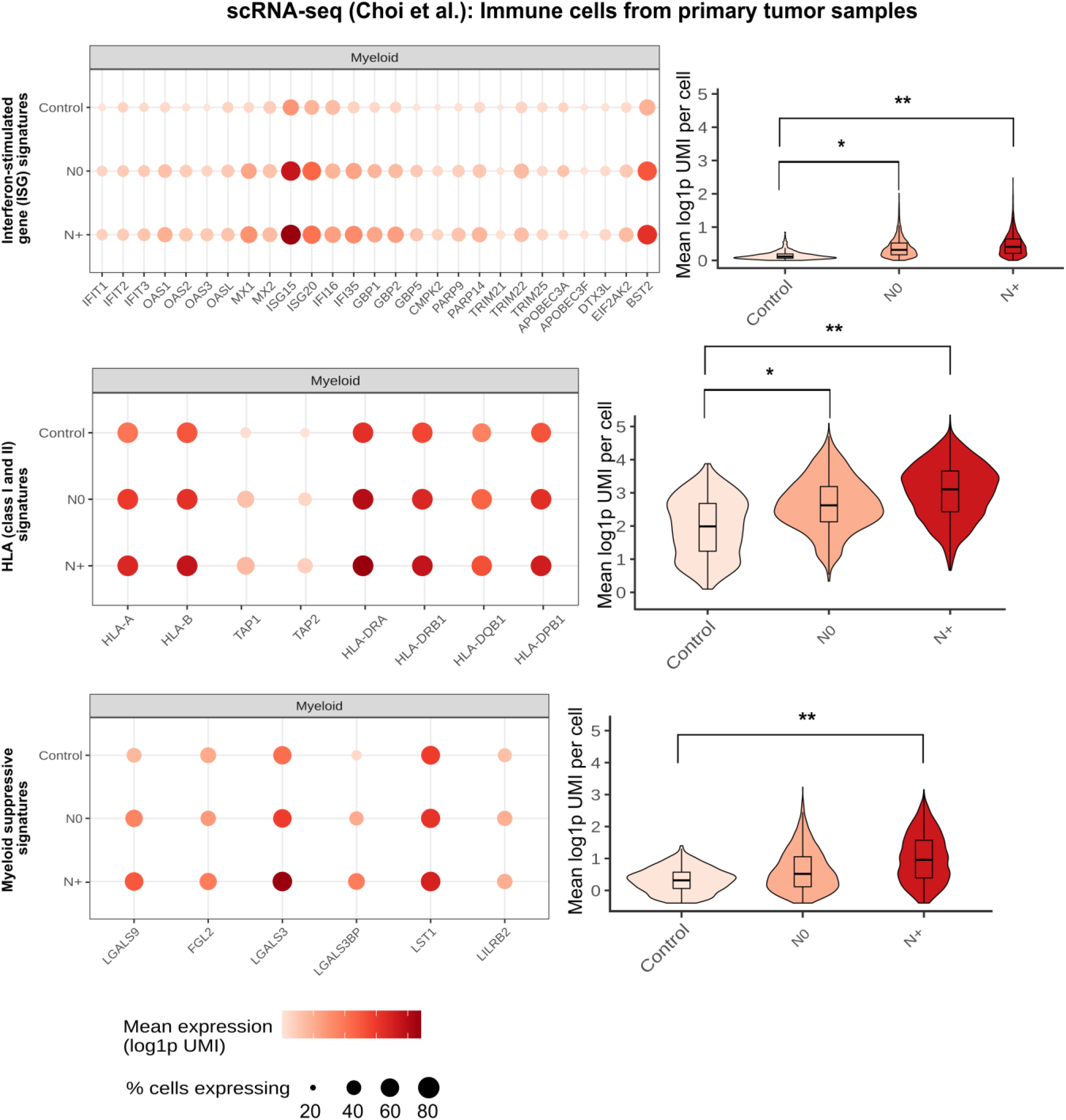
Independent scRNA-seq analysis reveals a convergent interferon-associated myeloid program. Analysis of the independent scRNA-seq dataset generated by Choi et al. (n=19 patients) of myeloid cells isolated from primary HNSCC tumors. Dot plots and violin plots illustrate the expression of ISGs, MHC, and myeloid suppressive signatures in myeloid populations from controls, pN0, and pN+ groups. Brackets indicate significant comparisons, with * *adjusted-p* < 0.05, \*\**adjusted-p* < 0.01, \*\**adjusted-p* < 0.001. Comparisons not reaching statistical significance are not shown.

Together, these observations provide orthogonal evidence for a conserved interferon-associated myeloid program across molecular layers and biological compartments. The scRNA-seq analysis independently identified an interferon-responsive myeloid program associated with clinical disease state, whereas SCP directly quantified the corresponding protein landscape in circulating immune cells. Thus, the two datasets provide complementary rather than directly comparable measurements, with SCP adding protein-level resolution of immune-cell states, signaling-associated proteins, and effector molecules.

### Cytotoxic lymphocyte programs become progressively dysregulated during metastatic progression

To determine whether metastatic progression was also associated with remodeling of protein coordination within circulating lymphocytes, pairwise correlations among proteins involved in T-cell receptor (TCR) signaling, adhesion, cytotoxic effector function, degranulation, and cell-cycle progression in PTPRCAP⁺ cells and its CD8^+^ subset were performed (**Figure 7**). The abundance of these proteins was subsequently evaluated according to the clinical groups and Leiden clusters from Figure 2 analysis (**Figure 7**).

**Figure 7.**
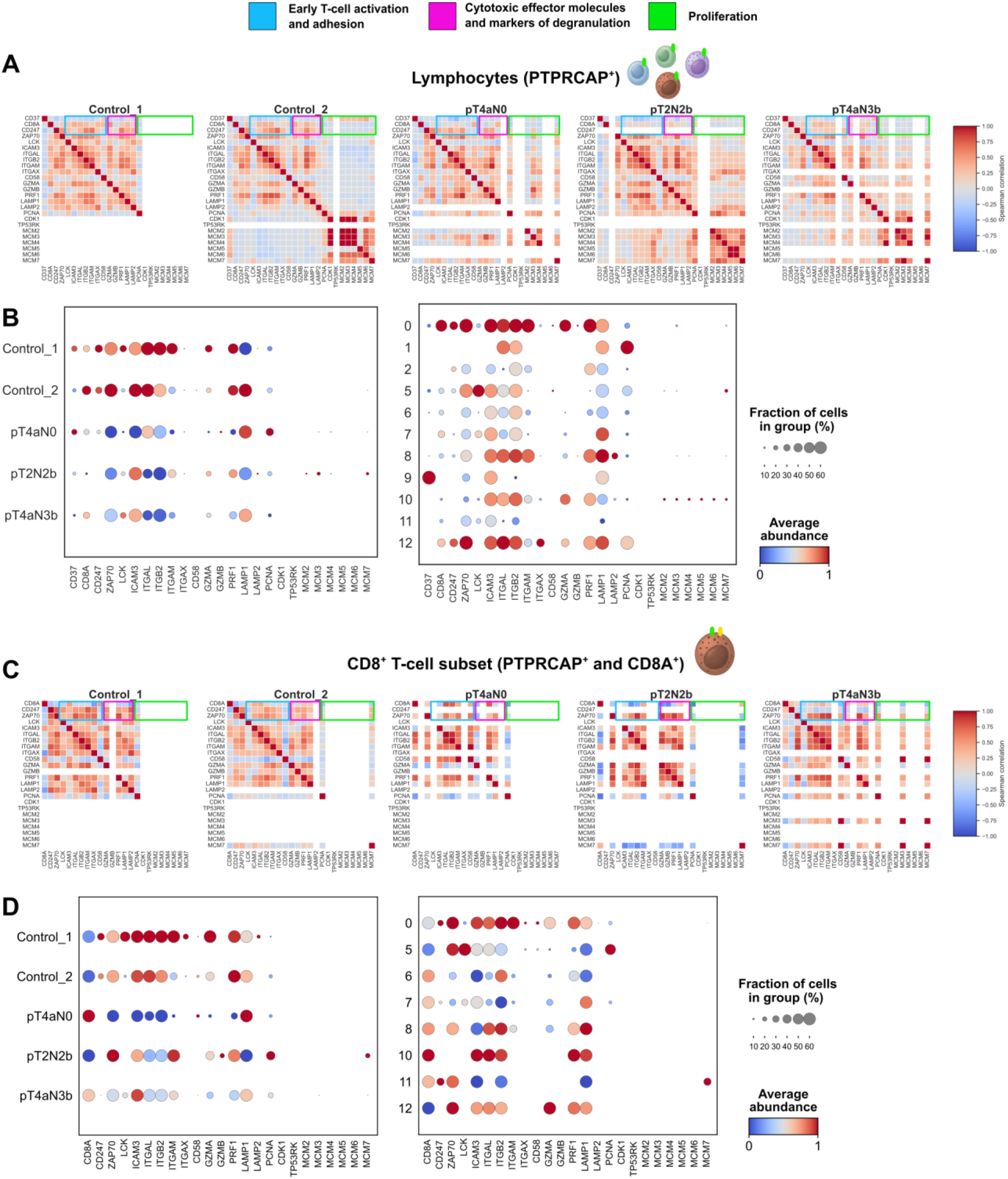
Coordinated remodeling of activation, cytotoxic, and proliferative protein programs in circulating lymphocytes during HNSCC progression. **(A)** Spearman correlation matrices showing coordinated protein abundance across circulating lymphocytes (PTPRCAP⁺ cells; n=253) from each clinical group (Control_1, Control_2, pT4aN0, pT2N2b, and pT4aN3b). Proteins were grouped according to their functional roles in early T-cell activation and adhesion (blue), cytotoxic effector molecules and degranulation markers (magenta), and proliferation (green). Correlation coefficients range from −1 (blue) to +1 (red). **(B)** Dot plots showing the abundance of the same proteins across clinical groups (left) and Leiden clusters (right) within the lymphocyte population. Dot size represents the fraction of cells expressing each protein, and color indicates the scaled average protein abundance. **(C)** Spearman correlation matrices for the CD8⁺ T-cell subset (PTPRCAP⁺ and CD8A⁺ cells; n=67) across the five clinical groups. The same functional protein categories shown in **(A)** are highlighted. **(D)** Dot plots showing protein abundance across clinical groups (left) and Leiden clusters (right) within the CD8⁺ T-cell subset. Dot size indicates the fraction of cells expressing each protein, whereas color represents the scaled average abundance.

In the overall lymphocyte population, controls displayed coordinated co-abundance among proteins involved in early T-cell activation, adhesion, and cytotoxic effector function ^40–43^. In contrast, the correlation patterns became progressively reorganized across HNSCC patients, with the loss of several coordinated associations and the emergence of new ones, indicating extensive remodeling of the lymphocyte proteomic network (**Figure 7A**). The accompanying dot plots also showed that proliferation-associated proteins were restricted to a limited subset of lymphocytes (cluster 10), which was not classified in a specific immune cell population (**Figure 2**). Besides, proteins involved in activation and cytotoxic function were distributed across multiple lymphocyte clusters with variable abundance; however, HNSCC patients presented a lower protein abundance distribution compared to both controls (**Figure 7B**). Interestingly, cluster 5, previously classified as CD4⁺ T cells and with higher abundance of cells from the pN+ patient with the more advanced stage of metastasis (**Figure 2**), exhibited the highest LCK abundance (**Figure 7B**), consistent with the central role of this kinase in TCR signaling and T-cell activation ^41^.

Next, to determine whether these changes were also evident within cytotoxic lymphocytes, we explored this same subset of proteins, except CD37 (cluster 9, specific for B-cells), in the cells with CD8A protein detected. Similar remodeling of protein co-abundance networks was observed, with progressive reorganization of activation- and cytotoxic-associated correlations across disease stages together with the emergence of a coordinated proliferative module in metastatic patients (**Figure 7C**). Proliferation-associated proteins were less prominently represented within the CD8⁺ T-cell subset and remained confined to a small fraction of cells, whereas activation- and cytotoxic-associated proteins were broadly distributed across multiple clusters (**Figure 7D**). Nevertheless, when focusing on the patient groups, a pattern similar to that observed in the overall lymphocyte population was observed, with HNSCC patients exhibiting lower average abundance of activation- and cytotoxic-associated proteins than both controls.

Collectively, these findings suggest the presence of two coordinated immune remodeling programs during metastatic progression. While circulating monocytes exhibited an interferon-associated inflammatory and immunosuppressive phenotype, cytotoxic lymphocytes displayed progressive reorganization of activation- and effector-associated protein networks.

Although the functional consequences of these coordinated changes remain to be established, warranting further investigation in larger patient cohorts, they point to a systemic immune landscape characterized by concurrent myeloid activation and altered cytotoxic lymphocyte organization more pronounced in the pN+ samples explored in the present study (**Figure 8**).

**Figure 8.**
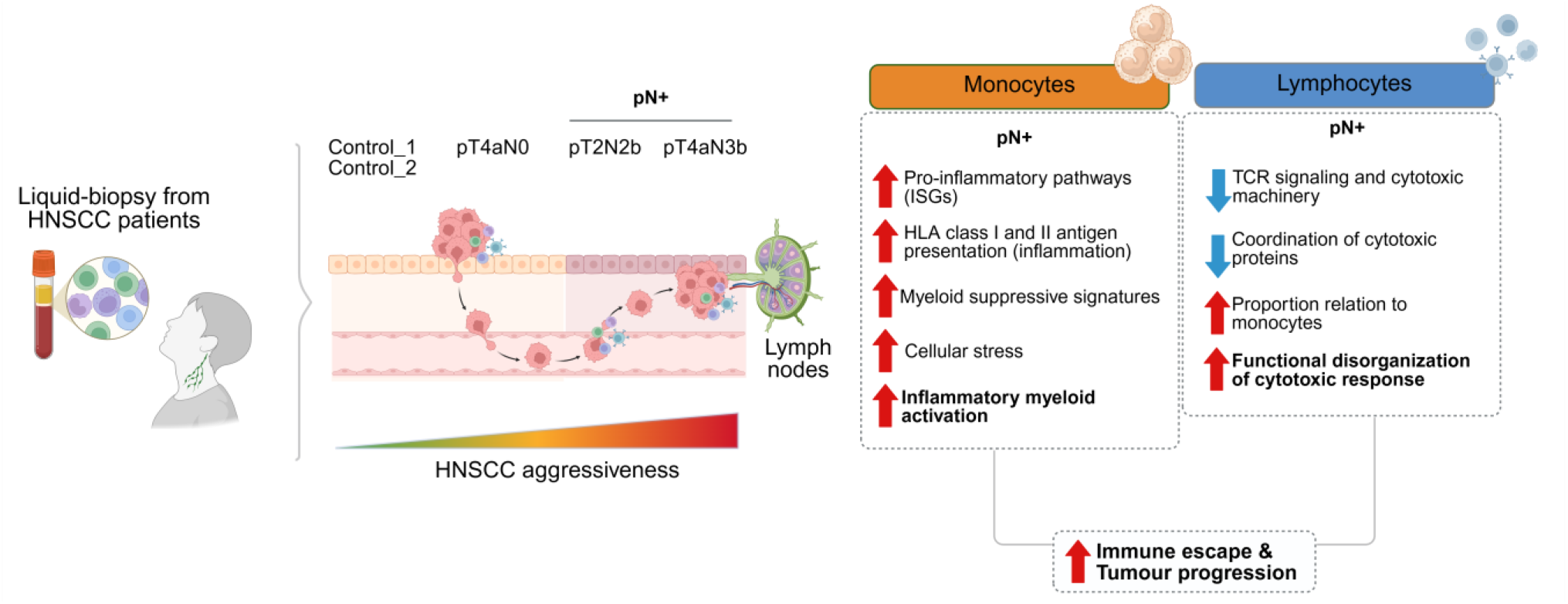
Exploratory biological interpretation hypothesis based on the SCP data from HNSCC patients and controls. As disease progresses to nodal metastasis, circulating monocytes acquire a progressively enhanced inflammatory and interferon-responsive phenotype, characterized by increased abundance of proteins associated with interferon signaling, antigen presentation, migration, and immunosuppressive myeloid functions. In contrast, cytotoxic lymphocytes exhibit progressive disruption of coordinated functional programs, with reduced coordination among proteins involved in T-cell receptor (TCR) signaling, adhesion, cytotoxic effector function, degranulation, and proliferation, accompanied by increased expression of selected inflammatory mediators. Together, these coordinated remodeling programs suggest a systemic environment that favors immune dysfunction, contributing to immune escape and metastatic progression.

## Discussion

Despite major advances in SCP over the past decade, relatively few studies have successfully translated this technology using clinical samples (liquid and tissue biopsies) ^13,14,17^. Technical challenges, including limited sample availability, low throughput, the need for standardized preservation strategies, and robust analytical workflows, have hindered its implementation in translational research ^17^. In this study, these challenges were overcome by applying a label-free SCP workflow, previously benchmarked using cell line models ^44,45^, to cryopreserved PBMCs from patients with HNSCC at distinct stages of nodal progression. Overall, we generated proteomic profiles of 619 single circulating immune cells and identified major immune populations present in peripheral blood by correlating Leiden cluster markers with bulk sorting proteomics data from Rieckman et al. ^23^.

Beyond immune cell-type identification, one of the most important findings of this study was the proteomic remodeling of both monocyte and lymphocyte compartments in advanced HNSCC states. These patterns were consistently observed in comparison with the two independent controls evaluated. Notably, these transitions were more pronounced in monocytes, suggesting that circulating myeloid cells are key drivers of systemic immune remodeling during HNSCC progression. We also observed a progressive reduction in the proportion of monocytes relative to lymphocytes in patients with HNSCC, indicating that disease progression is accompanied by both compositional and functional remodeling of the circulating immune compartment ^28,29^. The convergence of independent analytical approaches further supported the biological relevance of this trajectory. Specifically, the overlap between pseudotime markers and DAPs, analyzed using each control as an independent reference, revealed a shared group of ten proteins (BLVRB, CD36, GNAI2, LAP3, LGALS9, NHERF1, TMX1, PRAM1, TYMP, and VASP).

Rather than representing biomarkers, these proteins define complementary biological processes that collectively characterize monocyte functional remodeling during disease progression. For instance, LGALS9 is a well-established immunomodulatory galectin expressed by monocytes and macrophages that regulates inflammatory responses and promotes T-cell dysfunction through TIM-3 signaling ^33,34^, while CD36 participates in lipid uptake, scavenger receptor signaling, and inflammatory macrophage activation ^46^. Together, these proteins support the acquisition of an activated and immunoregulatory monocyte phenotype along the trajectory. A second group of proteins is associated with cell migration and dynamic cytoskeletal remodeling. GNAI2 regulates chemokine receptor signaling and leukocyte chemotaxis ^47,48^, whereas VASP controls actin polymerization, adhesion, and cell motility, processes required for immune-cell trafficking and interactions with the vascular endothelium ^49^. These coordinated changes suggest that monocytes progressively acquire molecular features associated with increased migratory and tissue-remodeling capacity.

Additional proteins point to metabolic and vesicle-associated adaptations. BLVRB and TMX1 contribute to redox homeostasis and protection against oxidative stress ^50,51^, whereas LAP3 participates in intracellular protein turnover and antigen processing ^52^. TYMP has dual functions in nucleotide metabolism and pro-angiogenic inflammatory signaling ^53^, while NHERF1 regulates membrane protein organization and intracellular signaling complexes ^54^. Collectively, these proteins are consistent with the enrichment of vesicle-mediated transport, intracellular trafficking, and immune regulatory pathways identified among trajectory-associated proteins.

The convergence of these independent functional modules supports the interpretation that the trajectory analysis captures a coordinated program of monocyte proteome remodeling rather than computational ordering of heterogeneous cells. Moreover, the fact that these proteins were independently identified both as trajectory-associated features and as reproducible DAPs across the two control references argues that this molecular program is robust to the choice of trajectory root and reflects progressive immune adaptation accompanying metastatic HNSCC. Nevertheless, these findings should be validated in larger cohorts before broader biological conclusions can be drawn.

In the context of cancer biology, monocyte heterogeneity and interferon-driven immune programs have emerged as key regulators of tumor-immune interactions. Recent studies have demonstrated that inflammatory monocytes orchestrate antitumor T-cell responses through interferon-dependent mechanisms while simultaneously undergoing functional reprogramming in chronically inflamed tumor environments ^55,56^. Interestingly, our SCP data identified cluster 3, characterized as monocyte cells exclusive of pN+ groups, as the immune population most closely associated with disease progression. These cells exhibited a broad interferon-response program characterized by increased abundance of canonical interferon-stimulated proteins together with enhanced MHC class I and II antigen-presentation machinery. The concomitant induction of interferon-stimulated proteins and antigen-presentation molecules suggests sustained interferon signaling within this monocyte population.

Although the present proteomic data do not allow us to discriminate between type I and type II interferon responses, the coordinated induction of canonical interferon-stimulated proteins together with MHC class I and II antigen-presentation machinery supports sustained interferon signaling within this monocyte population. The coordinated upregulation of canonical interferon-stimulated proteins together with HLA class I antigen-processing machinery (HLA-A, HLA-B, TAP1/2, TAPBP, and ERAP1/2) is consistent with activation of an interferon-responsive antigen-presentation program in circulating monocytes.

In this context, interferon-responsive monocytes simultaneously upregulated antigen-presentation machinery and immunoregulatory proteins, including LILRB2 and LGALS1/LGALS3, suggesting the acquisition of a phenotype in which enhanced antigen presentation coexists with immunoregulatory features during metastatic progression ^57^. Such dual activation states have been described in inflammatory myeloid populations exposed to persistent interferon signaling and may contribute to immune dysregulation within the tumor microenvironment ^58,59^. Nevertheless, further functional studies will be required to determine whether this interferon-responsive monocyte population primarily supports antitumor immunity or instead promotes immune dysfunction during metastatic progression. Resolving this balance may have important translational implications by informing therapeutic strategies targeting interferon signaling or immunoregulatory myeloid programs.

Interestingly, monocyte subclustering revealed heterogeneous distributions across clinical groups and distinct monocyte-associated programs. Monocyte subclusters enriched in pN+ patients showed a greater tendency toward intermediate and nonclassical monocyte-associated programs. This observation is consistent with the recognized heterogeneity of circulating monocytes, in which classical, intermediate, and nonclassical subsets exhibit distinct inflammatory, antigen-presentation, endothelial-interaction, and tissue-trafficking profiles ^37,39^. Nevertheless, these results should be interpreted as exploratory rather than definitive phenotypic classifications, and the observed pattern in the SCP data supports the hypothesis that metastatic HNSCC is associated with functional differences in circulating monocytes. Some canonical proteins used to define these monocyte subsets, such as CD16 (FCGR3A), a key marker of nonclassical monocytes, were not detected in our SCP dataset, limiting the ability to assign definitive phenotypic identities to single monocytes. This limitation underscores the need to further optimize SCP workflows for the efficient recovery and detection of membrane-associated proteins, which are essential for resolving closely related immune cell phenotypes at the single-cell level.

Beyond the myeloid compartment, our SCP dataset also uncovered potential functional alterations within circulating cytotoxic lymphocytes. Correlation analyses revealed progressive loss of coordination between canonical T-cell receptor signaling proteins, including LCK and ZAP70, and cytotoxic effectors such as PRF1 in metastatic patients. Similar patterns were observed in CD8⁺ lymphocytes. These observations are consistent with previous scRNA-seq studies reporting exhaustion-associated transcriptional programs in tumor-infiltrating CD8⁺ T cells and NK cells from patients with nodal metastasis ^11,12,60^. Our results extend these findings by demonstrating that systemic functional dysregulation of cytotoxic lymphocytes can also be detected directly at the proteomic level in circulating PBMCs.

Overall, our findings support an exploratory model in which metastatic progression in HNSCC is characterized by coordinated but distinct remodeling of the circulating immune system. While monocytes progressively acquire an interferon-driven inflammatory and immunoregulatory phenotype, lymphocytes exhibit reduced coordination between activation and effector programs. This imbalance between interferon-activated myeloid cells and functionally dysregulated cytotoxic lymphocytes may contribute to ineffective antitumor immunity, facilitating immune escape and metastatic dissemination.

Finally, this study also demonstrates the feasibility of applying SCP to cryopreserved clinical PBMC samples without prior cell sorting, enabling an unprecedented proteome characterization of systemic immune states directly from peripheral blood. By integrating an independent transcriptomic dataset as an orthogonal molecular layer, our work demonstrates how SCP and scRNA-seq provide complementary views of immune-state remodeling. The scRNA-seq data capture transcriptional programs within tumor-associated myeloid cells, whereas SCP directly measures protein abundance and resolves protein-level cellular states in circulating immune cells.

Together, these findings highlight SCP as a promising platform for resolving systemic immune states and its potential evaluation for longitudinal immune monitoring and biomarker discovery in larger cohorts.

## Resource availability

All data are available in the main text or supplementary materials. The mass spectrometry raw files were deposited at PRIDE under the code access: PXD080834. The code generated in this work will be made publicly available in a GitHub repository upon publication of the manuscript: https://github.com/HeloisaMonteiroAP/SCP_Paper_PBMC.

## Acknowledgments

We are grateful to the patients who donated blood to this research project, to the hospital team who collected the samples, to the drivers who transported the samples from the hospital to the laboratory, and to Dr. Fernanda Salvato from Thermo Scientific and Dr. Marvin Thielert for fruitful discussions on improving the LC-MS/MS and cellenONE X1 methods, respectively. We also extend the acknowledgements to The Mass Spectrometry Laboratory staff (Proposal number MAS - 20252312) and the Biobank, Premium Network, Instituto do Câncer do Estado de São Paulo - ICESP, Faculdade de Medicina, Universidade de São Paulo - USP, São Paulo, SP 01246-000, Brazil. Figures 1 and 8 contain illustrations from the BioRender repository (https://www.biorender.com/).

## Author contributions

Conceptualization: HMAP, AFPL

Methodology: HMAP, JGM, NARC, AFBL, DF, RRD, BAP, ALM, ACPRS, TB, LPK, AFPL

Investigation: HMAP, AFPL Visualization: HMAP, NC, JGM Funding acquisition: AFPL

Project administration: HMAP, AFPL Supervision: AFPL

Writing—original draft: HMAP

Writing—review and editing: HMAP, JGM, NC, TM, RNR, AFBL, AFPL

## Declaration of interests

The authors declare they have no competing interests.

## Declaration of generative AI and AI-assisted technologies

During the preparation of this work, the authors used ChatGPT (version 5.3) to improve the readability of some parts of the manuscript. After using this tool, the authors reviewed and edited the content as needed and take full responsibility for the content of the published article.

## Funding

This work was supported by Fundação de Amparo à Pesquisa do Estado de São Paulo (FAPESP) grants EMU 22/11476-5 and 2018/18496-6 (AFPL), and scholarships 23/12076-3 (HMAP), 23/16823-8 (JGM), 25/04523-5 (NARC), 22/12815-8 (DF). Conselho Nacional de Desenvolvimento Científico e Tecnológico (CNPq) grant 310392/2021-7 (AFPL). This work was also supported by or used resources from the Brazilian Federal Government that were provided to the Brazilian Center for Research in Energy and Materials (CNPEM), a private nonprofit organization under the supervision of the Brazilian Ministry for Science, Technology, and Innovation (MCTI). This study was conducted at the Mass Spectrometry Laboratory of the Brazilian Biosciences National Laboratory (LNBio), which is part of CNPEM.

## Materials and Methods

### Ethical approvals and blood collection

Blood samples were collected from healthy individuals and patients with head and neck squamous cell carcinoma (HNSCC). The study was approved by the Ethics Committee of the Piracicaba Dental School (FOP), São Paulo, SP, Brazil, through protocol CAAE: 54995522.0.0000.5418. Blood was collected at the Instituto do Câncer do Estado de São Paulo, located in the city of São Paulo and the state of São Paulo, Brazil.

### Patient and Control Cohort

Peripheral blood samples were obtained from two healthy control individuals and three patients diagnosed with HNSCC. Clinical characteristics of the cohort are summarized below. The control group included a female donor (47 years old) (Control 1) and a male donor (79 years old) (Control 2). The HNSCC cohort consisted of patients presenting distinct pathological stages and primary tumor sites, including tongue and retromolar trigone tumors. One patient presented localized disease without nodal involvement (pT4aN0 – 62 years old), whereas the remaining patients exhibited advanced nodal metastatic disease (pT2N2b – 59 years old, and pT4aN3b – 43 years old).

### Cell culture of HeLa cells

HeLa cells (RRID:CVCL_0030) were purchased from the Rio de Janeiro Cell Collection (Banco de Células do Rio de Janeiro, Brazil) and were authenticated by the ATCC. The cells were maintained in Dulbecco’s modified Eagle medium supplemented with 10% fetal bovine serum (FBS) (Cultilab, F063) and 1% antibiotics (penicillin and streptomycin) (Gibco, 15140-122) at 37°C and 5% CO2. For single-cell proteomics (SCP) analysis, cells were cultured in medium-sized flasks (T75) and collected at 80%–90% confluence. Next, cells were trypsinized (Cultilab, 517) and washed twice with 1× phosphate-buffered saline (PBS) (Gibco, 18912-014).

### Peripheral blood mononuclear cell (PBMC) extraction and cryopreservation for SCP

The collected peripheral blood was transported at room temperature from the hospital to a laboratory where it was quickly processed to extract and cryopreserve the PBMCs for subsequent SCP experiments. Cells were extracted from 16 mL of peripheral blood in EDTA tubes from healthy controls (n = 2) and patients with HNSCC (n = 3), diluted in PBS at a 1:1 proportion, and carefully added to the same volume of Ficoll®-Paque Plus density gradient media (Cytiva, 17144003) without mixing. Then, the solution was centrifuged at 600× *g* for 30 min and PBMCs were collected, followed by two washes with PBS and centrifugation at 200× *g* for 5 min.

Following the extraction, 1 × 10^7^ cells/mL of PBMCs were cryopreserved using a freezing medium composed of 90% FBS and 10% DMSO. Cells were rapidly transferred to a freezing container (Mr. Frosty, Sigma, C1562) for 16 h and stored in an ultra-freezer (−80°C). PBMCs from patients with HNSCC and controls were stored at −80°C once received at the hospital until the day of the SCP experiments (more than 8 days of storage at −80°C). After the cryopreservation period, cells were rapidly thawed in a dry bath at 37°C, washed twice with PBS, followed by 200× g for 5 min to remove the freezing media, and stained for viability prior to single-cell isolation.

### Cryopreservation of HeLa cells used as quality control (QC) for SCP

HeLa cells (1 × 10^6^ cells/mL) were cryopreserved using a freezing medium composed of 95% FBS and 5% dimethylsulfoxide (DMSO) (Sigma, 472301). Cells were rapidly transferred to a freezing container (Mr. Frosty, Sigma, C1562) for 16 h and stored in an ultra-freezer (−80°C). After the cryopreservation period, cells were rapidly thawed in a dry bath at 37°C, washed twice with PBS to remove the freezing media, and stained for viability prior to single-cell isolation.

### Detection and isolation of cells assisted by the cellenONE X1 robot

Cell isolation and sample preparation for SCP were performed using a cellenONE X1 robot (Cellenion®, Lyon, France). All solutions were prepared using MS-grade reagents, degassed prior to use, and loaded into a medium-sized uncoated piezoelectric dispensing capillary (PDC) (Scienion, Bico Company, C00016). Before cell isolation, cells were incubated with a Zombie NIR™ fixable viability kit (BioLegend, 423106) at a final concentration of 1:250 for 15 min at room temperature. The cells were then washed twice with PBS and resuspended in filtered PBS (Millipore® Steritop® vacuum bottle top filter, Merck, SCGPS02RE) at a final concentration of 200–300 cells/µL.

A master mix, consisting of 100 mM triethylammonium bicarbonate buffer (TEAB) (Sigma, T7408), 0.05% n-dodecyl-β-D-maltoside (DDM) (Sigma, D4641) and 3 ng/µL trypsin (Waters, 186010108) in de-gassed MS-grade water, was dispensed into each well of a clean Eppendorf LoBind 384-well PCR plate (Merck, EP0030627300). The transmission channel was initially used for cell selection, followed by fluorescence channel analysis using a red LED for the negative selection of viable cells (T > F). To ensure analysis of all cell particles, the detection parameters were set to a wide range: diameter of 3–100 µm and elongation up to 4. The following cell isolation parameters were optimized for each cell population: HeLa cells (n = 10 for each plate), diameter of 20–40 µm and elongation up to 1.8; PBMCs, diameter of 9–30 µm and elongation up to 1.8. During the cell isolation, the 384-well plate was maintained at 10°C and 45% humidity. Next, the plate was incubated at 50°C for 2 h. The humidity was maintained at 85% to minimize evaporation, and wells were hydrated with 500 nL of water. In the second round of hydration, 500 nL of 3 ng/µL trypsin resuspended in de-gassed MS-grade water was used, as described by Matzinger *et al.* ^44^ and Bubis *et al.* ^45^. After 2 h of incubation, 4 µL of 0.1% formic acid (FA) (Thermo, LS118-212) was added to each well using an electronic repeater pipette to quench the trypsin activity. The plate was then dried in a Thermo SC250EXP SpeedVac concentrator (Thermo, SC250P1-115) at room temperature and stored at −20°C until injection. Wells containing multiple cells (n = 10 cells; denominated as a library) were processed through the same workflow to construct sample size-comparable spectral libraries ^61,62^.

In Table 1, we summarized the number of cells isolated in each control and clinical group.

**Table 1.** Clinical cohort and number of single cells analyzed.

| Group | Sample type | Clinical classification | TNM | Cells isolated | Cells included after QC |
| --- | --- | --- | --- | --- | --- |
| Control_1 | PBMC | Healthy donor | — | 98 | 95 |
| Control_2 | PBMC | Healthy donor | — | 133 | 131 |
| pT4aN0 | PBMC | HNSCC | pT4aN0 | 129 | 128 |
| pT2N2b | PBMC | HNSCC | pT2N2b | 134 | 132 |
| pT4aN3b | PBMC | HNSCC | pT4aN3b | 134 | 133 |
|  |  |  |  | **628** | **619** |

### Experimental controls for SCP

As experimental controls for SCP during cell isolation, we included blank wells on the cell isolation plates that contained only the master mix without any cells (n = 3 wells for each plate of PBMCs from patients). An additional cleaning step of the PDC (referred to as “sterilization”) was performed before and after cell isolation, with three solutions used in sequence (0.5% sodium hypochlorite, 3% hydrogen peroxide, and 70% ethanol), followed by a “sci-clean” task and two flushes of the PDC with 250 nL of MS-grade water at a wash station. Prior to proteomic analysis, wells were inspected, and those containing more than one isolated cell or contaminated by debris were excluded from subsequent analysis.

As a quality control (QC) measure for sample preparation and to assess consistency across multiday SCP experiments, single cryopreserved HeLa cells (n = 10 per plate) were included on every plate. All experiments were performed using aliquots obtained from the same cryopreserved HeLa cell batch to ensure that comparisons between QCs were not influenced by batch-to-batch variability.

### LC-MS/MS analysis

Samples in the 384-well plates were resuspended in 4.0 µL of 0.1% FA, and 3.5 µL was analyzed using a Vanquish Neo UHPLC chromatographic system (RRID:SCR_026495) coupled with an Orbitrap Astral mass spectrometer (RRID:SCR_026205). The chromatographic separation of peptides was performed on a C18 Aurora Ultimate TS column (25 cm × 75 µm ID, 1.7 µm. The column oven (IonOpticks, AUR4-25075C18) temperature was 50°C. The system maximum pressure was 1,200 bar with a loading volume of 1 µL, with fast loading and wash options enabled. A gradient elution was conducted at a throughput of 50 samples per day using an active gradient. The percentage of buffer B (80% ACN in water with 0.1% FA) initially increased from 0% to 40% over 19.5 min at a nominal flow rate of 200– 450 nL/min, followed by a gradual increase to 99% buffer B from 19.5 to 22 min at a flow rate of 300 nL/min, which was maintained for an additional 5 min.

Single-cell-derived peptides were acquired in positive mode using the FAIMS Pro interface (Thermo, FMS02-10001), with a compensation voltage set to −48 V and a spray voltage of 1,800 V. Orbitrap MS1 spectra were acquired at a resolution of 240,000, with a scan range of 400–800 m/z, a normalized automatic gain control (AGC) target of 500%, and a maximum injection time of 100 ms. Data-independent acquisition (DIA) of MS2 spectra was performed on the Astral mass spectrometer using the same scan range, an AGC of 800%, and loop control set to 0.6 s per cycle, with varying isolation window widths and injection times. Precursor ion fragmentation was achieved using higher-energy collisional dissociation with a normalized collision energy of 27%. Data acquisition was performed for 24 min once elution began. Following data collection, the FAIMS voltage was set to −70 V during the washing and equilibration steps to minimize contaminant transmission to the mass spectrometer.

### Data analysis

For DIA-NN ^63^ DIA searches (version 2.5), an *in silico* spectral library was created using the UniProt human reference proteome (UP000005640, 42,547 entries, downloaded on June 2026), with the “contaminant” option selected and supplemented with the common Repository of FBS Proteins (cRFP; 199 entries) ^64^ to eliminate common mass spectrometry contaminants and protein derived from FBS, respectively. Protein groups (PGs) from the contaminant repositories were removed. Blank samples were independently analyzed with MBR enabled in DIA-NN.

The parameters used for DIA-NN software are described in Table 2.

**Table 2.** DIA-NN parameters.

|  | DIA-NN version 2.5 |
| --- | --- |
| Mode | Prediction from FASTA |
| Protease | Trypsin/P |
| Missed cleavages | 2 |
| Mass variable modifications | 1 |
| Contaminants | On |
| N-term M | On |
| Acetyl (N-term) | On |
| Peptide length range | 7–30 |
| Precursor charge range | 1–4 |
| Precursor m/z range | 400–800 |
| Fragment ion m/z range | 400–800 |
| FDR (%) | 1 |
| MS1 accuracy | 0 |
| MS2 accuracy | 0 |
| MBR | On |
| Unrelated runs | On |
| Protein inference | On |
| Scoring | Peptidoforms |
| Proteotypicity | Protein names |
| Machine learning | NNs (cross-validated) |
| Quantification strategy | QuantUMS (precision) |
| Cross-run normalization | RT-dependent |
| Library generation | IDs, RT and IM profiling |
| Speed peak filtering | Optimal results |
| Speed RT/IM filtering | Balanced |

### Data filtering and integration with cellenONE morphometric information

PG tables generated by DIA-NN (pg_matrix.tsv) were first filtered to remove contaminants. Missing values were replaced with zeros. Proteins found in the blank controls were excluded, and cells with less than a 1.35-fold change based on the blank average were removed from subsequent analyses.

The morphology data (diameter, elongation, and circularity) were obtained using cellenONE software and integrated with the proteomic tables by remapping the row (1–24) and column (1–16) information provided by cellenONE to the well positions in the 384-well plates, with rows A–P and columns 1–24.

### Post-analysis Metrics

Bar charts displaying the average and standard deviation (SD) of the PGs and peptides were generated using the *sns.barplot* function in the Seaborn package (v0.13.2) ^65^. A linear regression was performed to assess the relationship between cell morphology and the number of PGs identified. Data normality was evaluated using the Shapiro–Wilk test. Pearson’s correlation was applied to normally distributed data, and Spearman’s correlation was used otherwise. Positive and negative correlations were indicated by ρ > 0 and ρ < 0, respectively. Statistical significance was defined as *p* < 0.05. Plots were generated using the *regplot* function in the Seaborn package (v0.13.2).

### Subcellular localization analysis

The subcellular localization of the identified PGs in sPBMCs was annotated using the Human Protein Atlas (HPA) ^66^. Protein gene symbols were matched to the HPA subcellular localization database. Gene symbols containing isoform annotations (e.g., “(Isoform 2)”) were standardized by removing the parenthetical annotation prior to matching against the HPA database. Proteins annotated with multiple primary localizations were assigned to all reported compartments. Individual HPA annotations were further consolidated into nine major cellular compartments (Nucleus, Cytoplasm, Plasma membrane, Vesicular system, Mitochondria, Endoplasmic reticulum, Golgi apparatus, Cytoskeleton, and Cell division), while annotations not belonging to these categories were classified as “Others”. The relative abundance of proteins assigned to each compartment was calculated as the percentage of all annotated proteins and visualized as a pie chart.

### Dimensionality reduction, heatmap, and trajectory analyses

Dimensionality reduction was performed using principal component analysis (PCA) and uniform manifold approximation and projection (UMAP) implemented in the Scanpy package (RRID:SCR_018139) ^67^. MS1 intensity values were transformed into a log format using a natural logarithm with a pseudocount of 1 (log1p). PCA was then performed on the log-transformed data. Next, UMAP plots were generated from a k-nearest neighbor graph constructed using the first 20 principal components with Leiden clustering ^68^ across a range of resolution parameters (0 to 3). The optimal resolution was selected based on biological interpretability and marker expression patterns, prioritizing the identification of biologically meaningful cell populations while avoiding over-partitioning into clusters lacking distinct proteomic signatures.

Differential expression analysis was performed using the Wilcoxon rank-sum test implemented by the *sc.tl.rank_genes_groups()* function, in which cells in a Leiden cluster were compared with all other cells. Cluster markers were filtered using the following criteria: adjusted *p* < 0.05, log fold change > 1, abundance in > 60% of cells within the cluster (pct_in), and abundance in < 20% of cells outside the cluster (pct_out). Dot plots were generated to visualize the expression patterns of the abundant protein markers across clusters with per-protein standard scaling enabled (*standard_scale = “var*”). The filtered results were exported as tab-delimited files for downstream analyses.

Hierarchically clustered heatmaps were generated using *sns.clustermap()*. Marker expression was normalized per protein using the canonical gene denomination and z-score transformation. Cells were annotated according to Leiden clusters, cell morphology features (diameter, elongation, and circularity), and trajectory information. Euclidean distance and average linkage clustering were applied. All graphics were generated using Python 3 in a Google Collaboratory notebook environment.

Trajectory analysis was conducted using Monocle3 (RRID:SCR_018685) ^69^ with default parameters in R (v4.4.1). Genes associated with pseudotime were identified using a graph-based Moran’s I test and ranked by q-value, applying a false discovery rate (FDR) threshold of 1 × 10⁻⁴.

### Volcano plots

The terminology for conditions was standardized, and experimental controls were used as references. Zero values, corresponding to missing data, were treated as missing values (NA). For each clinical group, exploratory differential protein abundance was assessed at the single-cell level using Welch’s t test against the corresponding control group, and p-values were adjusted for multiple testing using the Benjamini–Hochberg false discovery rate (FDR) method. Log2 fold changes were calculated using a pseudocount of 1. Proteins with |log2 fold change| ≥ 1 and FDR-adjusted p < 0.05 were considered candidate differentially abundant proteins for exploratory interpretation. All plots were generated using Python 3 in a Colaboratory notebook environment

### Gene Ontology annotation

Gene Ontology (GO) enrichment analysis was performed using the g:Profiler Python interface (gprofiler) (RRID:SCR_006809) ^70,71^ to identify enriched biological processes and cellular components among differentially abundant proteins. Analyses were restricted to *Homo sapiens* and the GO Biological Process (GO:BP) category. Enriched terms were retained at a significance threshold of adjusted p-value < 0.05 after g:SCS multiple testing correction. Enrichment significance was expressed as −log10-transformed adjusted p-values. For each condition, the top 10 enriched GO terms ranked by adjusted p-value were selected.

### Immune cell population annotation based on the Human Gene Atlas and reference bulk-sorting proteomics of human immune cells

The cell type annotations were determined based on a search of the cluster markers against the BioGPS database available as a “Human Gene Atlas” option in the Enrichr web tool (RRID:SCR_001575) ^72,73^, This reference contains tissue- and cell type–enriched gene expression profiles that were used to identify the most likely immune populations. Cluster annotations were subsequently refined and validated using reference bulk-sorting mass spectrometry-based proteomics of purified human immune cell populations. ^23^.

Reference proteomes of 28 human hematopoietic cell types isolated from the peripheral blood of seven healthy donors were obtained from Rieckmann et al. and used to generate lineage-specific immune cell markers. These markers represented seven major immune populations, including granulocytes, monocytes, dendritic cells, natural killer cells, B cells, CD4⁺ T cells, and CD8⁺ T cells. For each immune population, protein abundances were compared with those of the remaining six lineages (Student’s t-test with Benjamini–Hochberg correction; adjusted p ≤ 0.05) to identify lineage-enriched proteins, which were further filtered to retain unique markers. SCP cluster markers were then compared with these reference lineage-specific protein signatures to assign immune cell identities to the clusters.

### Monocyte subset-associated protein scoring

To characterize monocyte subset-associated states, three independent protein signatures representing classical, intermediate, and nonclassical monocytes were defined based on established lineage-associated markers and information on highly abundant proteins reported for each monocyte subset in the Human Protein Atlas ^36–39^. The classical signature comprised CD14, S100A8, S100A9, FCN1, LYZ, VCAN, CTSZ, MNDA, and S100A10; the intermediate signature comprised HLA-DRA, HLA-DPA1, HLA-DPB1, CD74, CTSS, FCGR1A, LILRB1, LGALS3, and IFITM3; and the nonclassical signature comprised LRP1, LST1, LILRB1, SERPINA1, CEACAM8, and MPP8.

For each cell, a signature score was calculated independently for each monocyte subset using the *score_genes()* function implemented in Scanpy. This approach generates a score based on the expression/abundance of the selected signature proteins relative to a matched background of control features. The resulting values were stored as *Classical_score*, *Intermediate_score*, and *Non_classical_score*.

The resulting scores were subsequently compared within each monocyte subcluster using the paired Wilcoxon signed-rank test, since the three scores were calculated for the same cells. Comparisons included classical versus intermediate and classical versus nonclassical scores. Statistical significance was defined as *p-value* < 0.05.

### Single-cell RNA sequencing analysis

Single-cell RNA sequencing (scRNA-seq) data from Choi et al. ^11^ (N=19 patients, in which 6 are pN+; 7 are pN0, and 6 are Control) were processed using Seurat (v5) (RRID:SCR_007322) in R ^74^. Cells were retained after quality control filtering using the following thresholds: nFeature_RNA > 200; Feature_RNA < 7,500 and percent.mt < 20%. Cells failing these criteria were excluded. Genes expressed in fewer than 3 cells were removed. Data were normalized using Seurat’s NormalizeData function with scale.factor = 10,000. All downstream analyses were performed on log-normalized expression values (Seurat “data” layer).

### Clinical harmonization

Pathological nodal staging was harmonized across cohorts. N stage was extracted from clinical annotations and classified as N0, N1, N2, or N3. For selected analyses, stages N1–N3 were grouped and defined as N+.

### Cell-type annotation

Cells were assigned to immune subpopulations using curated marker-based annotation. Broad immune classes included monocytes and myeloid/dendritic cells (DCs). For statistical analyses, these classes collapsed into a single myeloid category comprising monocytes and macrophage/DC populations.

### Gene sets

Three predefined gene sets were evaluated: interferon-stimulated genes (ISGs), antigen presentation genes (HLA), and immunosuppressive genes. Gene set membership was curated a priori based on published literature.

### Pseudobulk aggregation strategy

All inferential analyses were performed using a pseudobulk framework to preserve patient-level independence and avoid pseudoreplication. For each gene g, expression values were aggregated across cells belonging to the same patient (p) and cell type (c) by computing the mean expression:

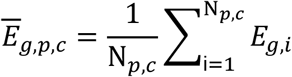

Where *E_g_*_,*i*_represents the expression of gene *g* in cell *i*, and *N_p_*_,*c*_ denotes the total number of cells from patient *p* within cell type *c*.

### Gene-set scoring

Gene-set scores were computed at the single-cell level as the arithmetic mean of the log-normalized expression values across all genes in each predefined gene set:

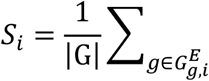

Where *S_i_*represents the gene-set score for cell *i*, *G*denotes the set of genes in the predefined gene set, ∣ *G* ∣is the number of genes in the set, and *E_g_*_,*i*_corresponds to the log-normalized expression value of gene *g*in cell *i*.

### Statistical analysis

Differential expression analyses between clinical groups were performed using one-way ANOVA followed by Tukey’s multiple-comparison post hoc test.

## Notes

### Competing Interest Statement

The authors have declared no competing interest.

